# Self-organization of conducting pathways explains electrical wave propagation in cardiac tissues with high fibrosis

**DOI:** 10.1101/455964

**Authors:** Nina Kudryashova, Aygul Nizamieva, Valeriya Tsvelaya, Alexander Panfilov, Konstantin Agladze

**Affiliations:** Department of Physics and Astronomy, Ghent University, Ghent, 9000, Belgium; Laboratory of Biophysics of Excitable Systems, Moscow Institute of Physics and Technology, Dolgoprudny, 141701, Russia; Laboratory of Computational Biology and Medicine, Ural Federal University, Ekaterinburg, Russia

## Abstract

Cardiac fibrosis occurs in many forms of heart disease and is considered to be one of the main arrhythmogenic factors. Regions with a high density of fibrosis are likely to cause blocks of wave propagation that give rise to dangerous cardiac arrhythmias. Therefore, studies of the wave propagation through these regions are very important, yet the precise mechanisms leading to arrhythmia formation in fibrotic cardiac tissue remain poorly understood. Particularly, it is not clear how wave propagation is organized at the cellular level, as experiments show that the regions with a high percentage of fibrosis (65-75%) are still conducting electrical signals, whereas geometric analysis of randomly distributed cells predicts connectivity loss at 40% at the most (percolation threshold). To address this question, we used a joint *in vitro-in silico* approach, which combined experiments in neonatal rat cardiac monolayers with morphological and electrophysiological computer simulations. We have shown that the main reason for sustainable wave propagation in highly fibrotic samples is the formation of a branching network of cardiomyocytes. We have successfully reproduced the morphology of conductive pathways in computer modelling, assuming that cardiomyocytes align their cytoskeletons to fuse into cardiac syncytium. The electrophysiological properties of the monolayers, such as conduction velocity, conduction blocks and wave fractionation, were reproduced as well. In a virtual cardiac tissue, we have also examined the wave propagation at the subcellular level, detected wavebreaks formation and its relation to the structure of fibrosis and, thus, analysed the processes leading to the onset of arrhythmias.

**Author summary:** Cardiac arrhythmias are one of the major causes of death in the industrialized world. The most dangerous ones are often caused by the blocks of propagation of electrical signals. One of the common factors that contribute to the likelihood of these blocks, is a condition called cardiac fibrosis. In fibrosis, excitable cardiac tissue is partially replaced with the inexcitable connective tissue. The precise mechanisms leading to arrhythmia formation in fibrotic cardiac tissue remain poorly understood. Therefore, it is important to study wave propagation in fibrosis from cellular to tissue level. In this paper, we study fibrosis of high density in experiments and computer simulations. We have observed a paradoxical ability of the tissue with extremely high fibrosis (up to 75% of fibroblasts) to conduct electrical signals and contract synchronously, whereas geometric analysis of randomly distributed cells predicted connectivity loss at 40% at the most. To explain this phenomenon, we have studied the patterns that cardiac cells form in the tissue and reproduced their self-organisation in a computer model. Our virtual model also took into account the polygonal shapes of the spreading cells and explained high arrhythmogenicity of fibrotic tissue.

## Introduction

The contraction of the heart is controlled by propagating waves of excitation. Abnormal regimes of the wave propagation may cause *cardiac arrhythmia,* asynchronous contractions of the heart and even lead to cardiac arrest and sudden cardiac death. Cardiac arrhythmias often originate from blocks of propagation [1]. In that case, the wave goes around the block, reenters the same inceptive region and, thus, forms a persistent rotational activity called *cardiac reentry,* which is one of the main mechanisms of lethal cardiac arrhythmias.

A normal heart has a complex structure, which is composed of the bundles of elongated cardiac cells. Apart from premier excitable cardiac cells, there are also inexcitable cells of connective tissue: *cardiac fibroblasts.* Their role is to maintain the structural integrity of the heart [2] and repair injuries [3]. Fibroblasts outnumber cardiomyocytes in a healthy human heart although occupying a much smaller total volume [2, 4]. However, many pathological conditions are associated with an excessive growth of the fibrous tissue, called *cardiac fibrosis,* which is, therefore, considered to be one of the major arrhythmogenic factors [5, 6].

The mechanisms of arrhythmia onset in fibrotic tissue remain poorly understood but generally believed to be associated with the increased probability of waveblock formation. It is a well-established fact that fibrosis slows down wave propagation [7] and can completely block it if the fibroblasts density is high. The critical density of fibrosis above which the conduction terminates is called *percolation threshold.* This concept originates from the percolation theory, and, by definition, it specifies the point of long-range connectivity loss/formation in random systems. Connectivity here refers to electrical synchronisation in the tissue, or, in other words, the ability to transmit electrical waves of excitation. Percolation threshold, i.e. the critical density of fibrosis which breaks long-range connectivity, plays an important role in arrhythmogenicity. It was shown that cardiac tissue is most susceptible to arrhythmias if the density of fibrosis is only slightly (~2-3% [8]) below the percolation threshold. There are two main factors that may facilitate reentry formation. First, a large amount of fibroblasts (acting as heterogeneities) increases the probability for waveblock formation [9]. Second, a high fraction of fibrosis creates a ‘maze’ that effectively lengthens the travel distance for the waves as they follow a longer zig-zag path [10] and, thus, provides sufficient room for the emerging reentrant loops. As a result, high density of fibrosis both facilitates the initiation and creates conditions for the existence of reentrant cycles, resulting in a highly arrhythmogenic substrate.

Up to now wave propagation failure was studied only in generic mathematical representations of cardiac fibrosis with each cell randomly chosen to be either myocyte or fibroblast (see e.g. [9, 11]). In this kind of 2D computer models, the propagation of excitation failed at 40% of fibrosis [12], which is also within the range of values predicted by classic mathematical models (e.g. 37-44% of the area uncovered by conducting elongated ellipses with the shape similar to cardiomyocytes [13]). However, experimental measurements [14] indicate that wave propagation and synchronous contraction in 2D cardiac monolayers is observed for up to 75% percentage of fibrosis.

In this paper, we study the phenomenon of wave propagation in cardiac tissue with a high density of fibrosis using a joint *in vitro-in silico* approach. We performed experiments in 25 monolayers with various percentages of fibroblasts and detected wave propagation to determine the percolation threshold. We have found that, indeed, the experimentally measured threshold (75% of the area covered by fibroblasts) is substantially higher than what was predicted in conventional computer modelling (40% [12]) or classic mathematical models. Further morphological examination revealed that the key mechanism of conduction in highly fibrotic tissue is likely to be tissue patterning. The cardiomyocytes were not located randomly but organised in a branching network that wired the whole sample.

Next, in order to explain cardiac network formation, we applied a virtual cardiac monolayer framework developed in [15], based on the Cellular Potts Models [16–18]. We proposed a hypothesis that such self-organisation occurs due to cytoskeletons’ alignment. Based on this hypothesis, we were able to obtain branching patterns, as well as reproduce the decrease in conduction velocity and wave percolation observed in experiments. We have further studied *in silico* the process of formation of the wavebreaks leading to reentry formation and analysed the tissue structures that caused them.

This paper is organized as follows. First, we describe the experiments conducted with the neonatal cell cultures. Second, we analyse the patterns and formulate the hypothesis on the mechanism of their formation. Third, we implement the hypothesis and reproduce pattern formation *in silico* in a Cellular Potts Model (CPM) of cardiac tissue [15]. Next, we compare electrical signal propagation in computer modelling and in experiments. Finally, we show a spontaneous formation of uni-directional blocks in the model resulting in reentry formation and locate and analyse the structures causing it.

## 1 Results

### 1.1 Experimental study of the percolation threshold in neonatal rat cardiac monolayers

We cultured neonatal rat ventricular cardiomyocytes mixed with cardiac fibroblasts in variable proportion (20-88% fibroblasts) and studied electrical activity in these monolayers using optical mapping.

In Figure 1a and in S1 Video the wave propagation in a sample with high fibrosis is shown (66% fibroblasts). In spite of high percentage of fibrosis, this sample is still conducting electrical waves, however the wave propagation pattern is complex. The wave originates from the stimulation point marked by a yellow spike at the bottom of the tissue and initially spreads in all directions. However, due to a large number of fibroblasts, the wave is blocked at multiple sites. After a short delay (in areas outlined with dashed ellipses), the wave propagates further into the left and the right parts of the sample, and again spreads in various directions. This process repeats multiple times, resulting in a complex, fractionated wave pattern containing narrow pathways and some bulk excited regions. The conduction blocks in Figure 1a are shown in red, which are the places where the wave propagation was blocked and the wave had to go around. The main propagation paths are shown with white and black arrows. In this sample, the electrical wave propagation was still possible and long-range connectivity was still present, however the amount of fibroblasts was close to the percolation threshold.

**Fig 1.**
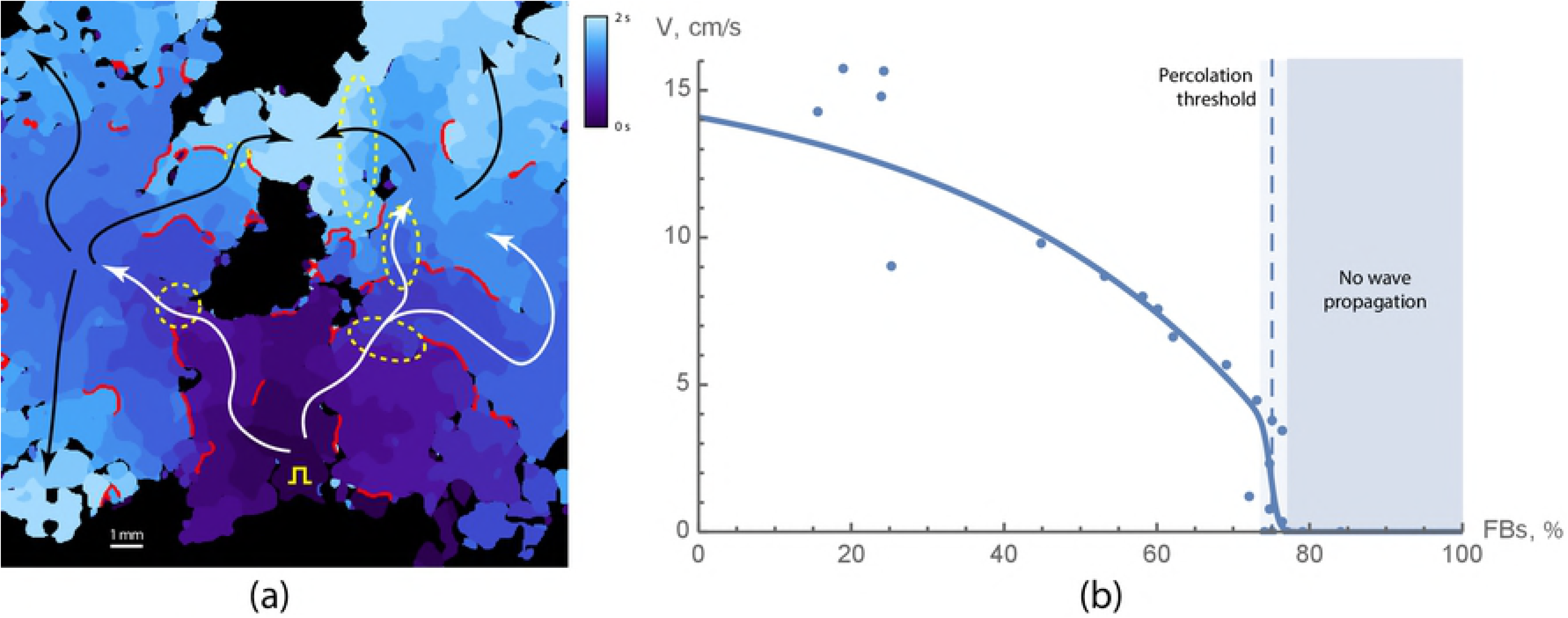
Wave propagation in a neonatal rat cardiac monolayers with fibrosis. (a) Activation map for a sample with 66% of fibroblasts. Activation times are colour coded. Red lines show the regions where the wave was blocked. White and black arrows show the main propagation pathways. Yellow square pulse sign indicates the location of the stimulating electrode. Yellow dashed lines outline the areas of slow conduction. The original video (S1 Video) of the wave propagation is available at https://youtu.be/3aDmsT1pl3Y. (b) Velocity decay with the increase of the portion of fibroblasts in samples. The percolation threshold is shown with the dashed line and was equal to 75 ± 2%.

We have found that the percolation threshold for the neonatal rat cardiac monolayers was 75 ± 2% of fibroblasts. We have also measured conduction velocities in the samples below the percolation threshold (Figure 1b). The measurements show that the velocity decreased when approaching the percolation threshold. In the samples with low fibrosis, the velocity was approximately 10-14 cm/s, and it decreased twofold in the samples with 70% fibrosis. The number of conduction blocks was higher in samples with high fibrosis. As a result, the mean conduction velocity decreased with the increase in a percentage of fibroblasts.

After optical mapping of the wave propagation, we have fixated the samples and studied their morphology using immunohistochemical labeling. We have found that the cardiomyocytes in the samples have formed connected networks capable of electrical wave propagation. In Figure 2 cardiomyocytes are shown in pink, and the cluster that they have formed is outlined with a white contour. The cardiomyocytes were organised in a branching structure that wired the whole sample. We have followed the pathway using a confocal microscope and it was possible to find long-range connectivity in the tissue. Therefore, we observed that the cardiomyocytes were organised in conduction pathways and assumed that there must be a mechanism responsible for their self-organisation.

**Fig 2.**
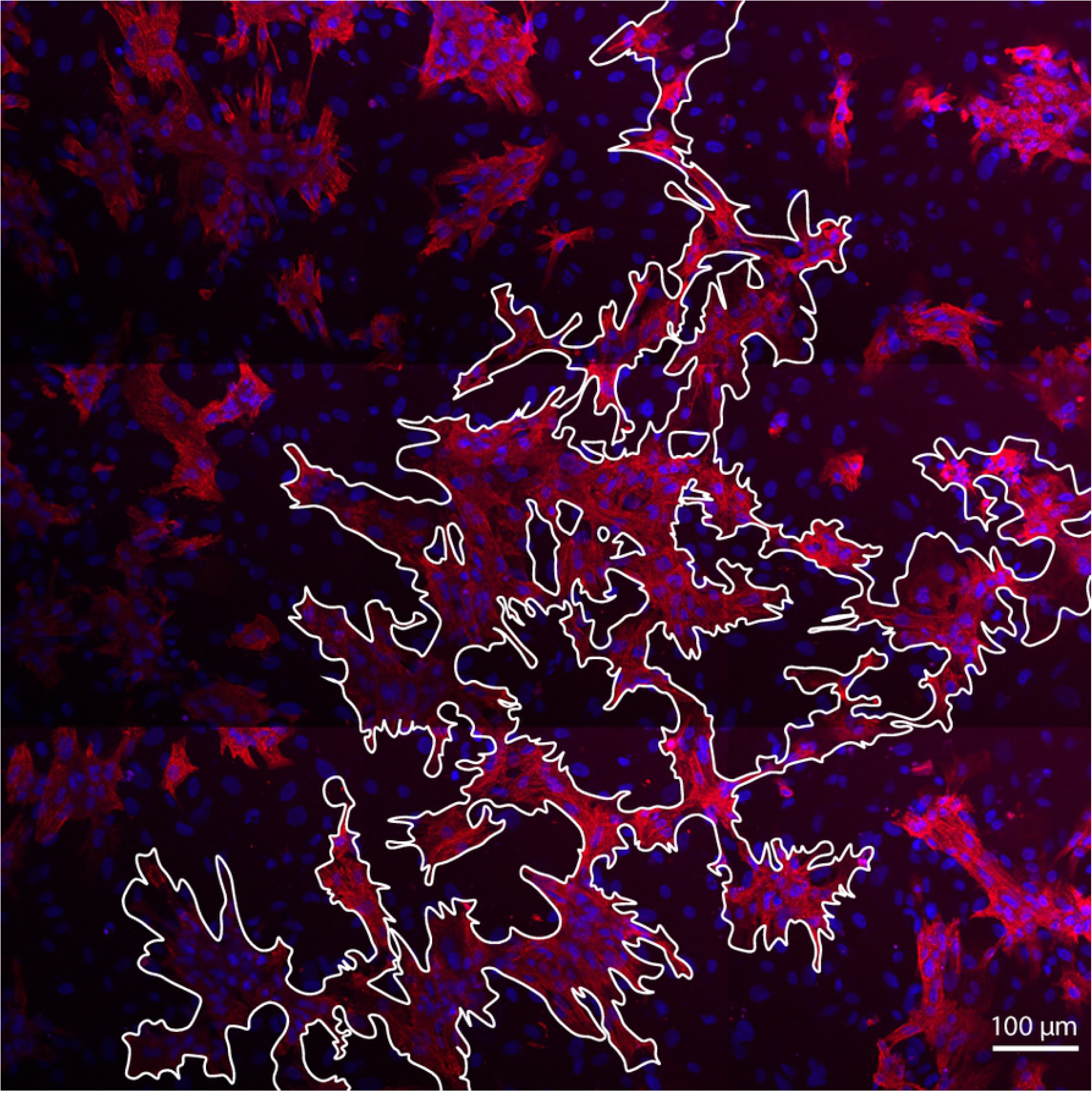
Conducting pathway in a monolayer of cardiac tissue with 31% of cardiomyocytes and 69% of fibroblasts. The interconnected region is outlined in white. Cardiomyocytes are labeled with anti-*α*-actinin antibody and coloured in pink. Nuclei are shown in blue.

### 1.2 Formation of branching structures in a computer model

Pattern formation in cell populations during development was extensively studied using Cellular Potts Models [16, 17, 19]. However, after trying several approaches used before (see the discussion section of our paper), we have not found an existing model that could have been applicable to the cardiac tissue and at the same time could reproduce the experimentally observed branching structures. Finally, after careful analysis of our experimental preparation, we found that an essential feature of our structure was the alignment of the cytoskeletons in the neighbouring cells. This alignment of the actin fibres is clearly seen in our experimental preparations. In Figure 3 the bundle of neonatal rat cardiomyocytes is shown. The red arrows indicate the intercalated disks between the cells. The white arrows point at the actin bundles on the opposite sides of the intercalated disk, that smoothly continue one another.

**Fig 3.**
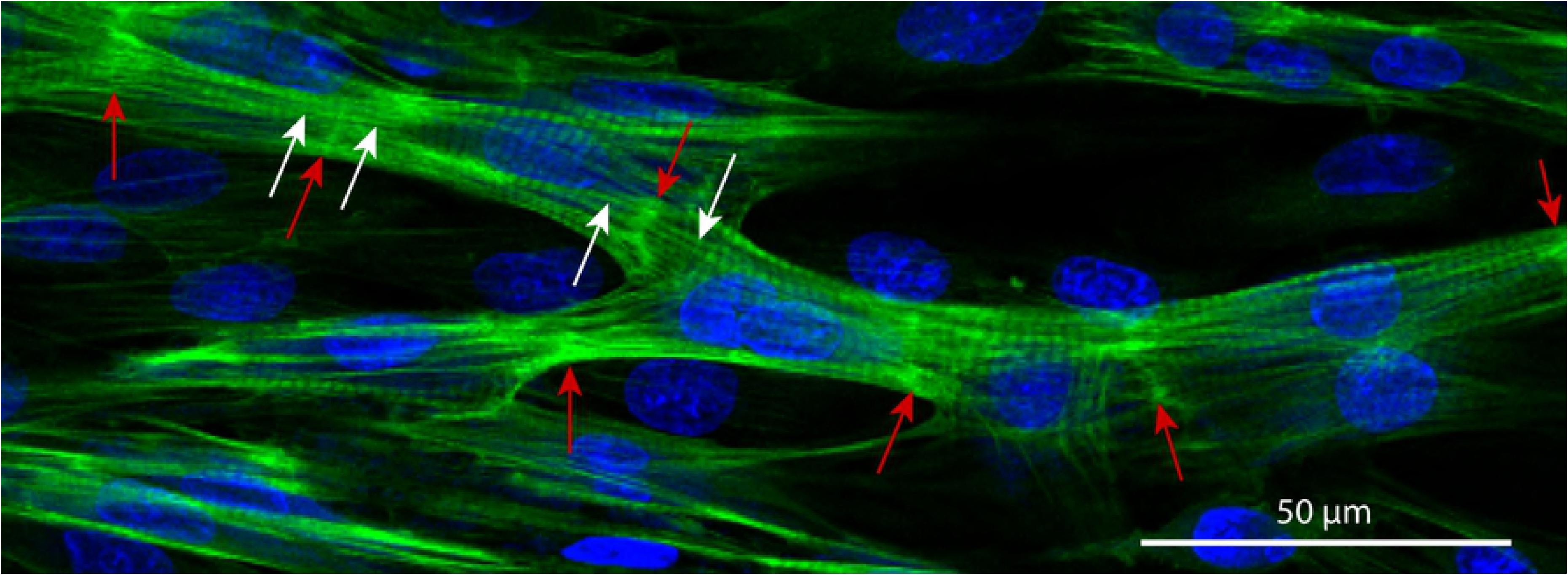
Cardiac syncytium in a 3-days culture of the neonatal rat cardiomyocytes. The cells have formed inercalated discs (ID, bright-green, highlighted with red arrows), aligned their cytoskeletons (the aligned strains on the both sides from ID are shown with white arrows) and formed a branching network. Nuclei are shown in blue (DAPI, labels DNA), and actin strands are shown in green (phalloidin, labels F-actin).

The precise biological mechanism of such alignment is unknown. However, in our view, it can be derived from a well-known property of actin cytoskeleton reinforcement in response to the external force [20]. Actin filaments, as well as the adaptors that link them, remodel under applied tension. In case of contact of two cells, the actin filaments are connected through adherens junctions, which transmit the tensile forces between the filaments of the neighbouring cells. It was shown that higher tension stabilises the whole complex [20]. The tension is maximal, if the actin filaments are aligned with each other, which gives a preference for intercellular alignment of the cytoskeletons.

Therefore, we have incorporated this mechanism into our Cellular Potts Model (CPM) of cardiac cells [15]. This model was already adjusted to reproduce characteristic shapes of the cardiac cells in virtual tissue model, and here we extended it with a new energy term, that corresponded to the alignment of actin bundles in the neighbouring cells.

In our CPM model, cell-substrate adhesion sites, to which actin bundles are anchored, were represented as separate entities: specially labeled subcells of the lattice. If two adhesion sites of two cells came into contact, we established the connection between them. The connection means, that the new bond energy was applied to them. This energy depended on the angle between the linked actin bundles (see Figure 4). The minimum of the energy corresponded to smooth coupling between the bundles, or zero angle. In this case, the bond was the most stable, but couplings with the non-zero angle between the bundles had a tendency to break apart.

**Fig 4.**
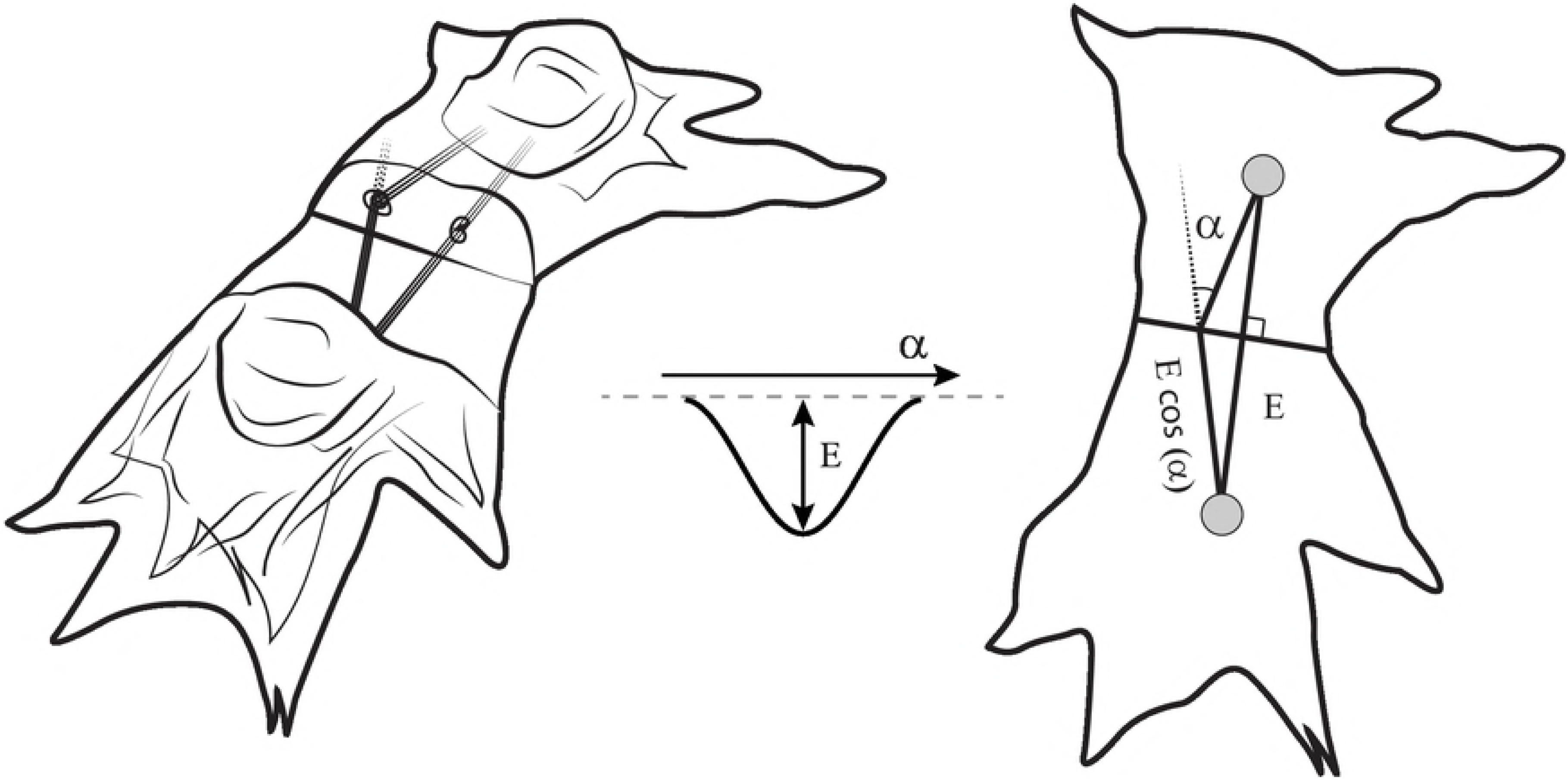
The schematic of the cell-to-cell interaction in a computational model. The energy term was assigned to every pair of connected actin bundles in coupled cells. This term depends on the angle between the bundles and reaches its minimal value when the bundles are aligned (*α* = 0). Left image shows quasi-3D schematic of the cells, middle image shows energy profile and right image shows a view from the top.

With this new energy term that favours cytoskeletons alignment, the cardiomyocytes in simulations created branching patterns. In Figure 5a a resulting simulated structure of the sample with 70% fibroblasts is shown. We see that in this sample only 30% of cardiomyocytes were able to build a network, and even with such a high density of fibroblasts, this network was fully interconnected. Our further studies showed that such interconnection was established for every sample with 30% of cardiomyocytes (*n* = 10) of 1 cm × 1 cm size, which is close to the size of the Petri dish used in our experimentals. For samples with 28% of cardiomyocytes the network was interconnected in 20% of cases. We also see that the patterns in computer simulation (Figures 5a) and in experiment (Figures 5b) have similar features, such as long single-cell-wide connections that bridged the gaps between cell clusters, or isolated clusters of fibroblasts trapped within the main cardiac pathway. Therefore, using the hypothesis on cytoskeletons’ alignment allowed us to reproduce the main features of the pattern.

**Fig 5.**
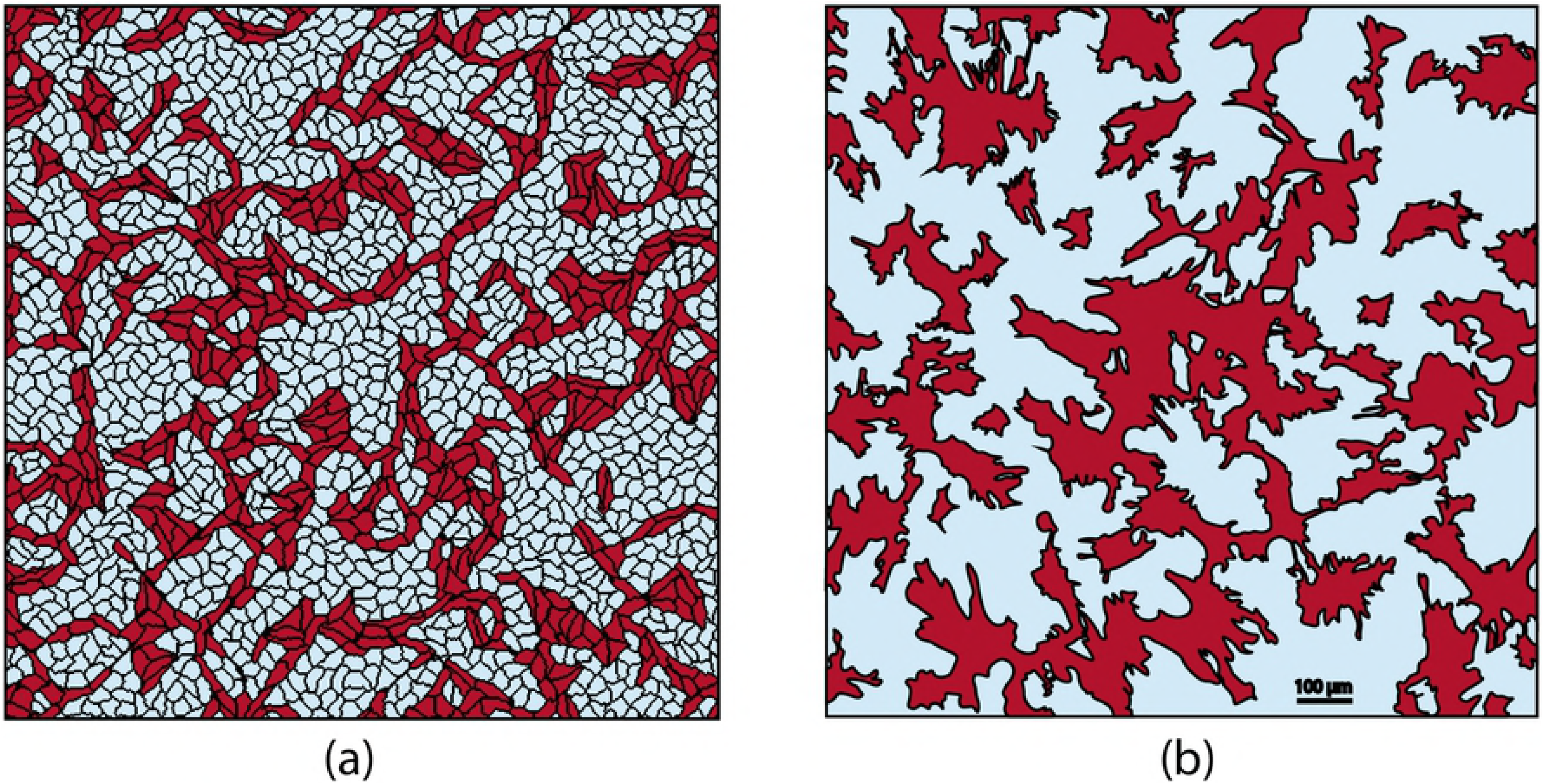
The branching pattern obtained in a computer model (a) compared to one observed in an experiment (b). **(a)** A virtual sample with 70% of fibroblasts. **(b)** A segmented image of the experimental sample with 66% of fibroblasts. The original image is shown in Figure 2.

S2 Video shows the growth of the branching pattern, highlighting the large connected clusters of the cardiomyocites by different colours. One can see from the video, that subtle movements of the cells (left) result in dramatic changes in connectivity (right). This emphasises the importance of the protrusions of the cells, which are taken into consideration in our model.

Using this model with cytoskeleton alignment, we have studied further wave excitation patterns and the percolation threshold for electrical waves in the system.

### 1.3 Wave propagation in virtual cardiac tissue monolayers

We have reproduced the experiments from Figure 1a *in silico* using Majumder et al. [21] model for neonatal rat ventricular cardiomyocytes.

Figure 6a and in S3 Video show the wave propagation in a virtual sample with 70% fibroblasts. The wave propagation pattern is complex and similar to one observed in experiment (see Figure 1a). One can see, that the number of percolation blocks per unit area is similar to that of the experimental activation pattern, and the trajectories of the waves share the same features. Note, that the scale of the simulated activation map is slightly smaller than those of the experimental one.

**Fig 6.**
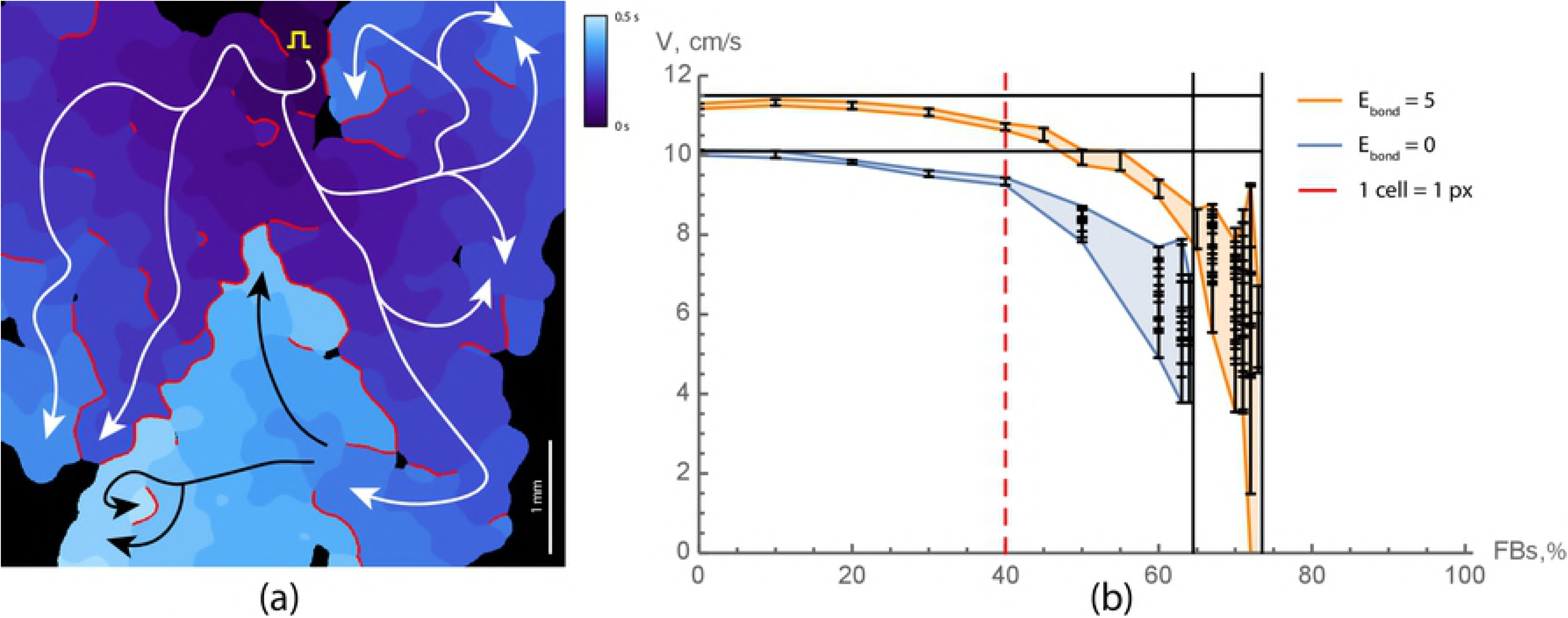
Wave propagation in virtual cardiac monolayers with fibrosis. **(a)** Activation map for a smaple with 70% of fibroblasts. Activation times are colour coded. Red lines show the regions where the wave was blocked. White and black arrows are showing the main propagation pathways. Yellow square pulse sign indicates the location of the stimulating electrode. **(b)** Velocity as a function of the fibrosis density. Orange lines represent velocities in virtual samples with cytoskeletons alignment (*E_bond_* = 5), and blue lines without (*E_bond_* = 0). Waves propagation in samples with branching patterns (orange) was failing in part of the samples starting from 71% and not possible in all of the samples for ≥75%. For each density, 10 samples with the size of 5 mm × 5 mm were tested. For each sample the mean velocity and its standard deviation are shown (mean ± SD). The original video (S3 Video) of the wave propagation is available at https://youtu.be/elvOvBRwnEM.

The percolation threshold in virtual cardiac monolayers was equal to 71.5 ± 1.5% of fibroblasts, meaning that 100% of the samples (*n* = 10) with 70% fibrosis were interconnected, whereas samples with more than 73% fibrosis (*n* = 10) were never conducting. For 72% fibrosis 20% of the samples were functional and for 71% fibrosis, those were 80%. The conduction velocity dependence on the density of fibrosis is shown in Figure 6b. For each simulated sample we have indicated the mean value of the velocity, and the standard deviation of the velocity distribution was shown with error bars. One can see, that the dependency of the velocity on the fibroblas density is similar to one measured in experiments (see Figure 1b). Closer to the percolation threshold the fluctuations in conduction velocities amplified due to the stochastic nature of the percolation block. Moreover, variations in conduction velocities within one sample were very high, because of a large number of conduction blocks.

The percolation threshold in our simulations was slightly lower (≈ 72%) than in experiment (75%). However, both values are much higher than the predictions of the models with random cells distribution (40%).

### 1.4 The unidirectional block and spontaneous reentry onset observed in virtual cardiac monolayers with high fibrosis

We have shown that in virtual tissues the spontaneous onset of reentry can occur. It was observed in samples with a high level of fibrosis. Figure 7 shows such process for a sample with 70% fibrosis. We see that after the first stimulus applied to the left border of the sample (shown in yellow), the wave propagation was blocked at the lower part of the sample, but continued propagation through the upper part. After reaching the right border, the wave turned around and entered the bottom region. However, this first wave was blocked in the middle of the sample, as the tissue in the upper part of the sample did not recover. The wave from the second stimulus, applied at the same site, has followed the same path and formed a sustained circulation along the path shown with the red dashed line in Figure 7.

**Fig 7.**
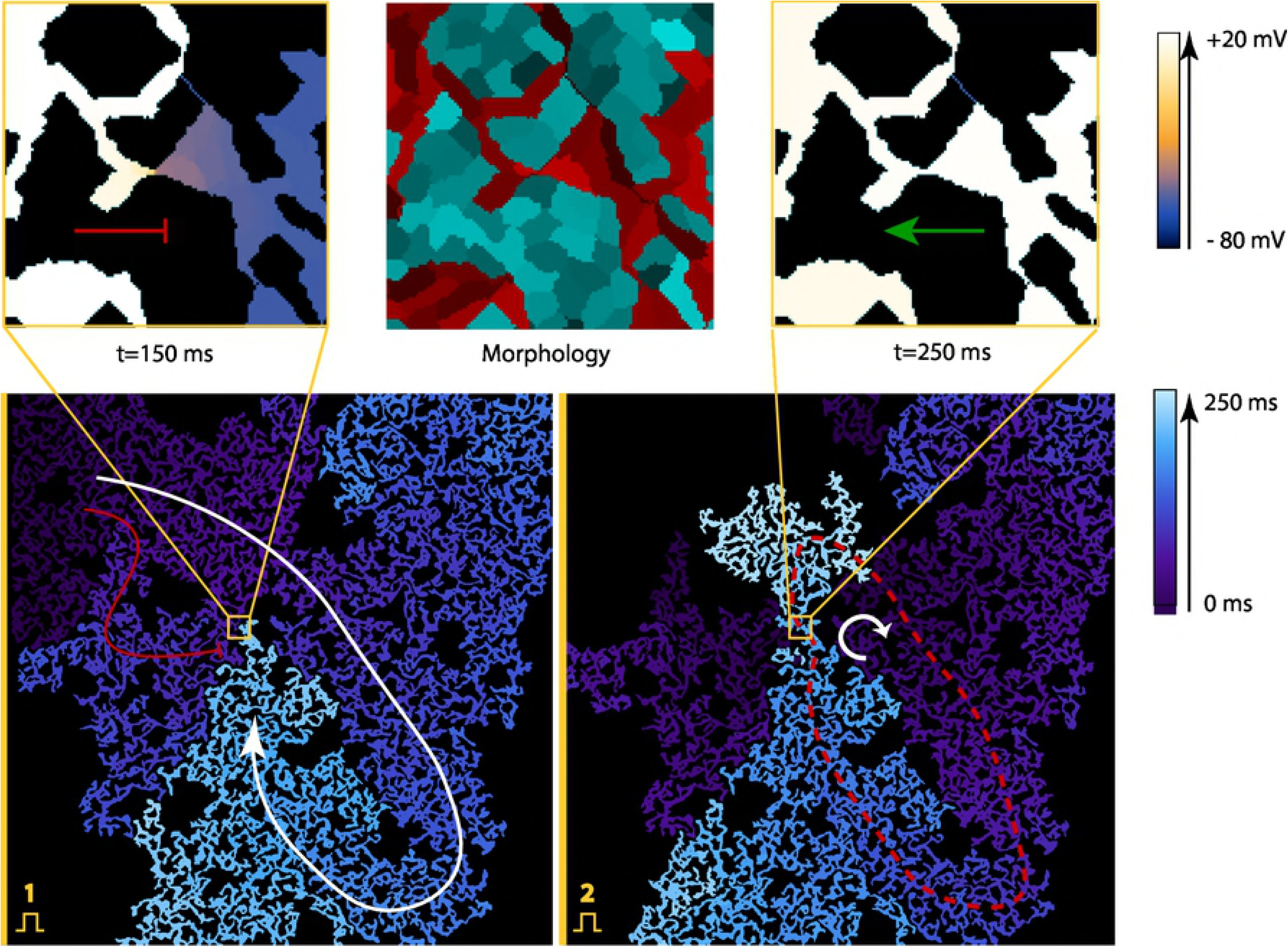
Spontaneous formation of the structure, that produces a uni-directional block, which results in reentry formation. The virtual sample had 29% of cardiomyocytes. Images on top show the insets of the place, where a uni-directional block has occurred: the central image shows its morphology and images on the side show wave propagation in different directions. The voltage on the top images is colour coded. In the central image, cardiomyocytes are represented with red tints and fibroblasts with cyan. The bottom images show the whole sample. The bottom-left image shows the uni-directional block after the first stimulus, which was applied on the left boundary. The bottom-right image shows the reentry formation after the second stimulus. The activation times are colour coded. The arrows show the main wave paths. The red dashed line represents the reentry cycle. The original video (S4 Video) of the wave propagation is available at https://youtu.be/6LZorTUcJdk.

Detailed analysis of the structure revealed that formation of a reentry here is solely due to specific structure which is shown in the middle (inside the yellow square) and acts as an area of uni-directional block (or “diode”): the waves can propagate from right to left, but not in the opposite direction. The diode is formed by two cell clusters that barely touch each other. These cell clusters have slightly different areas adjacent to this connection. Therefore, if the wave propagates from left to right (from a smaller cluster to a larger one), then the small cell does not produce enough current and can not depolarise a bigger one. The transmembrane potential in the largest cluster raises (which is shown in red colour), but not enough for the sodium channels to open. This effect is called *source-sink mismatch* [1]. When the source (a smaller cluster) is insufficient compared to the sink (a larger inactive cell cluster), the wave propagation is blocked. This effect was observed in chemical systems [22] and later in cardiac monolayers [23].

In a sample in Figure 7 the “diode” may initiate a reentry if the sample is stimulated from the left or from the top with a high frequency (4 Hz or more).

We have performed studies in 6 large samples with 70% of fibroblasts. All of these samples had 2-5 areas of the uni-directional block and many bi-directional blocks, but only one of these samples was arrhythmogenic. Thus, in addition to “diodes”, some extra geometrical conditions are required. The precise conditions are to be studied in the future, however, they are related to the presence of the long conducting circuits in the tissue. In fact, in Figure 7 we see that apart from the diode, there is also a loop, which is shown with a red dashed curve in the bottom-right image. The diode is a part of this loop, however, the loop is large enough to account for recovery of the bottom part of the tissue after one rotation. The samples with a higher density of fibrosis were even less likely to have long circuits, thus we didn’t observe sustained reentry there. We concluded, that reentry formation requires not only uni-directional blocks but also long circuits, and the intermediate densities of fibrosis are the most arrhythmogenic ones. This result was previously shown for random cell distributions, that reentry is most likely to occur 5-10% below the percolation threshold [8]. The principle holds true in our model, but quantitatively the densities of fibrosis are different.

Our model shows, that the areas of the uni-directional block can be naturally formed during tissue growth. Every time one spreading cell comes in contact with the other cell cluster, there is a chance that this connection will be asymmetric. It may explain the fact that reentries are frequently observed in the experimental setups with cardiac monolayers.

## 2 Discussion

We observed paradoxical electrical wave propagation in samples with up to 73–75% of fibroblasts instead of mathematically predicted 40% for randomly distributed cells. We have shown both *in vitro* and *in silico,* that electrical wave propagation was possible due to the formation of the conduction pathways that rewired the whole monolayer. We have proved the existence of this branching network with immunohistochemical images. We have measured the conduction velocity, which decreased with the increase of the portion of fibroblasts in the monolayer in a similar way in experimental and computational studies. Assuming that cardiomyocytes align their cytoskeletons to fuse into cardiac syncytium, the morphology of conductive pathways was successfully reproduced in computer modeling as well as the electrophysiological properties of the monolayers.

To explain the formation of the pathways, we have considered several hypotheses. First, we suggested that differential adhesion together with cell elongation may be enough to explain this patterning. A similar system with autonomously elongating cells was successfully used to explain vasculogenesis [24]. However, we did not impose obligatory elongation on the cardiac cells, because they are not necessarily elongated according to our experimental observations. Cardiomyocytes obtain their typical brick-like shape with the guidance of the extracellular matrix over the course of development. However, there is no evidence for any internal autonomous mechanism for elongation. In experiments on the glass, cells were not only bipolar but also tripolar and multipolar. In our model, this feature was reproduced with explicit introduction of the actin bundles. Similar approach was previously used to describe the shapes of dendric cells [25]. Cooperation between aligned actin bundles pulling in one direction shaped the cell more efficiently, which resulted in clustering of these bundles and multipolar cell shapes. Therefore, since the elongation was not imposed, there was no mechanism forcing cardiomyocytes out of clusters to search for new connections.

Next, we considered various mechanisms that were previously used to explain similar branching patterns in angiogenesis. The main sources of instability in those models were chemotaxis and contact-inhibition [26]. The percolation in the networks formed due to sharp gradients of chemoattractants was also studied previously in a continuous model [27]. However, there was no evidence for directed migration of cardiac cells and for type-specific contact-inhibition, similar to those that select a tip cell in a growing blood vessel. Therefore, we discarded this hypothesis either.

This mechanism is similar to *diffusion limited aggregation:* a process in which random-walking particles form fractal aggregates. In our model, a cell does not only stick to the growing pattern but also aligns its orientation with it. The mechanism for it actually exists as a part of syncytium formation, when cardiomyocytes fuse and align their cytoskeletons. Once a cell aligns cytoskeleton with its neighbours, this structure maturates and fixes the cell in place. In some sense, this process causes contact-inhibition only between cardiomyocytes and not between cardiomyocytes and fibroblasts, which results in the branching structure like one shown in angiogenesis [26].

Formation of conduction pathways and complex texture of the tissue may be important not only in terms of arrhythmogenicity. For example, it was shown that texture of cardiac tissue at subcellular level can substantially affect the propagation of external current during defibrillation [28, 29]. It would also be interesting to quantify the excitation patterns in terms of the number of re-entrant sources and wavefront complexity [30]. Therefore, it will be interesting to perform a similar study for the textures generated with our model.

There are several limitations of the methods used in this study. First of all, we have conducted experiments with cell cultures, which are different from the real cardiac tissue. It would be interesting, yet more complicated, to study patterning of the real 3D cardiac tissue. However, the mechanisms of patterning that we discovered here could be universal and might be applicable to the real cardiac development as well. Second, fibrosis is a more complex condition than just excessive growth of fibroblasts, which also involves collagen deposition. The collagen electrically insulates the cardiac fibers from one another and is also considered to increase arrhythmogenicity. In this study, we did not take the extracellular matrix into account, but it would be also interesting to measure its effect on the percolation threshold in the future studies. Finally, percolation depends on the size of the sample and scales with this size. For random systems, the scaling laws are known. In our case, we did not consider scaling and used similar size of the samples *in silico* as in the experiment. However, it would be interesting to study scaling of the percolation threshold and see how the size can affect the probability of wavebreak formation.

We conclude, that the cardiomyocytes in fibrotic areas can form a connected network and allow electrical signal propagation in monolayers containing up to 75% of connective tissue.

## 3 Methods

### 3.1 Experimental samples preparation

#### Neonatal cardiac cell isolation

All studies conformed to the Guide for the Care and Use of Laboratory Animals, published by the United States National Institutes of Health (Publication No. 85-23, revised 1996) and approved by the Moscow Institute of Physics and Technology Life Science Center Provisional Animal Care and Research Procedures Committee, Protocol #A2-2012-09-02. In this study, we used enzymes adapted to the existing two-day protocol selection from Worthngton-Biochem^1^. Cardiac cells were isolated from the ventricles of rat pups (Rattus norvegicus, Sprague Dawley breed) with different ages (1–4 days). Then, the isolated cells were seeded on the specimens covered with fibronectin (0,16 mg/ml, Gibco, USA, 33016015) at different concentrations before they were cultivated in DMEM culture medium (Gibco, USA, 11960) with 5% of FBS (foetal bovine serum, Gibco, USA, 10100147). For the study of the shapes of the isolated cells, the cells were seeded at 5 · 10^3^ cells/cm^2^. After 3 days of cultivation, the samples were fixated. The monolayers of primary culture cells were seeded at 30 · 10^3^ cells/cm^2^, and after 3–5 days monolayers were used in morphometrical studies and optical mapping.

#### Immunohistochemical staining

The cells were fixated with 5% PFA (paraformaldehyde powder, 95%, Sigma-Aldrich, USA, 158127-100G), and nuclear staining was performed with DAPI (VECTASHIELD Mounting Medium with DAPI, Vector, USA, Cat. No. H-1200). In our work, we used anti-*α*-actinin (Sigma-Aldrich, USA, A7811)and alexa fluor 594 like Secondary Antibody (A-11020, Life Technologies) for CM-specific labelling, Alexa Fluor 488 phalloidin (Molecular Probes, USA, A12379) for F-actin non-specific staining and DAPI for labelling cell DNA. Pictures were taken with an inverted fluorescence microscope (Axio Imager with ApoTome optical sectioning module, Zeiss). Immunofluorescent staining of the CMs was performed with the use of secondary and primary antibodies according to a previously described protocol^2^. Signals from each fluorescent label were recorded in a corresponding wavelength range. Channels were then preudo-coloured and merged together.

#### Optical mapping

To monitor activity and record the excitation patterns, the 3- to 5-day-old monolayers were loaded with the Ca^2+^-sensitive indicator Fluo-4-AM (Molecular Probes, USA, F14201). After staining, the medium was exchanged with Tyrode’s solution (Sigma-Aldrich Co., USA, T2145-10L) and kept at room temperature during the observations. The excitation waves were monitored with a high-speed imaging setup (Olympus MVX-10 Macro-View fluorescent microscope equipped with high-speed Andor EM-CCD Camera 897-U at 68 fps). All videos were processed with ImageJ software.

#### Velocity measurements

To measure velocity of excitation wave propagation, especially for samples with low concentration of CMs and thereby fluctuating velocity values, 10 space intervals for each sample were selected. The velocity was measured for each interval and after the mean velocity and standard deviation were calculated.

#### Cell counting

Two techniques were applied for cell counting.After immunostaining, cells were counted either manually or by means of cytofluorometry. For manual counting cells’ nuclei stained with DAPI represented the overall number of cells, while only those within *α*-actinin stained fibers were accounted for cardiomyocytes. The second method was performed on cytofluorometer (Accuri C6, BD Biosciences, USA) by distinguishing non-cardiomyocytes, stained only for F-actin, from CMs, stained both for F-actin and *α*-actinin

#### Mathematical model for cardiac tissue growth

In this study, we have used a mathematical model that was previously developed for cardiac monolayers formation [15]. It is based on the Cellular Potts Model (CPM) formalism, which describes cells as the domains of the regular lattice and assigns energy to the system of cells to describe their growth and motility. In our previous work [15], we have selected the main features of the cardiac cells (such as area, number of protrusions, etc.) and parametrised our model to reproduce the cell shapes. In this study, we extended the model with a new energy term responsible for syncytium formation.

The evolution of the cardiac monolayer is described by the Hamiltonian:

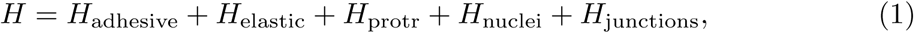

where *H*_adhesive_ + *H*_elastic_ is the basic CPM model and *H*_protr_ is the term describing the protrusion dynamics of the cardiac cells, which produces a characteristic polygonal shapes of these cells. *H*_nuclei_ corresponds to higher rigidity of the nuclei compared to the cell body. Finally, *H*_junctions_ is a new term, that describes the stability of adherens junctions and alignment of the cytoskeletons of the neighbouring cells.

The core feature of the model of cardiac cells is the explicit representation of cell attachments as the labels on the subcells of the lattice. They are first assigned when the cell expands, and can be destroyed with a certain penalty, if, for example, the cell is stretched due to its movement. The number of attachments per cell is limited by the amount of actin present in the cytoplasm, which is also reflected in the model. If the maximal number of attachments is reached, the new attachments do not form.

The spreading of virtual cells is shown in S5 Video. One of the extra features of our approach, is that elongation of the cells is not imposed. Cardiac cells in a real heart are indeed elongated, as they are guided by the extracellular matrix, whereas in a petri dish cells can spread in any direction with no preference. However, this does not result in a circular shape due to the limited number of attachment sites where spreading occures. Therefore, instead of explicit declaration of elongation, we suggested that virtual cells have only a small (10) number of well-developed mature attachment sites. Moreover, these sites in a model tend to cluster stontaneously (see S5 Video), as the actin strands pull the cell more efficiently acting together, rather than separately. Therefore, these clustered attachment sites turn cell shapes into bipolar, tripolar or multipolar states. The bipolar state is the most stable one for isolated cells, however tripolar cells are also relatively common. This correctly represents the population of cells observed in experiments.

*H*_protr_ is an energy term, that decreases with the distance from the centre of mass of the cell, and which is applied only to these labeled attachment sites. As a result, the attachment sites spread out and reproduce a characteristic polygonal shape of the cardiac cells. Here, in this model, we omit the details of the spreading process, which involves polymerization and depolymerisation of the actin, attachment/detachment, etc., but we mimic the overall dynamics of the protrusions. Also, we assume, that for every attachment site a corresponding actin bundle exists, which stretches from the attachment site towards the proximity of the nuclei.

If two attachment sites in the neighbouring cells come into contact, the bond between these attachment sites can be formed. In the algorithm, this bond appears if one attachment site attempts to move over the other. In such case, instead of copying of the subcell the connection establishes. A new energy term *H*_junctions_ applies to the subcells, which are involved in a newly established junction. This term determines the stability of the cell-to-cell junction and depends on the angle between the actin bundles, associated with the attachment sites involved (see Figure 4). It is determined as follows:

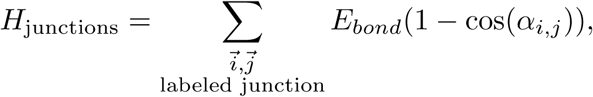

where *α_i,j_* (also shown in Figure 4) is the angle between two cytoskeleton bundles of two neighbouring cells which ends at points *i* and *j* which are labeled as the parts of the junction. The higher is the angle, the less stable is the junction. Therefore, the junctions with continuous actin bundles on both sides persist, but the kinked bundles tend to lose the connection. As a result, the actin bundles of the neighbouring cells tend to align and stay in the aligned state.

The parameters of the model used in simulations (see Table 1) in this paper were adjusted to compensate the additional energy term *H*_junctions._ The most of them are close to the parameters used in our original paper [15]. The value of the new energy term was set to *E_bond_* = 5.0, which provided enough stability for the junctions to maintain the branching structure but at the same time not too much stability to allow cells to search for possible new connections. Addition of *H*_junctions_ effectively increased the adhesion between cardiomyocytes. Therefore, the differential adhesion was toned down in a model to allow cardiomyocytes to migrate randomly before they maturate and stick to the pattern. In this study we have changed type-specific adhesion coefficients *J_cell−cell._* Our choice of the parameters was guided by the desired balance between branching and clustering: 1) lower adhesion energy (J) to the cells of the same type increased the size of unstructured, irregularly shaped clusters; 2) on the other hand, high adhesion energy forced cells to migrate out of clusters. We have chosen neutral values of *J_CM−CM_* = *J_CM−FB_*, which meant that cardiomyocytes had no preference in neighbours. Their energy was equal for being surrounded with either other cardiomyocytes, or with the fibroblasts. The fibroblasts had a slight tendency to cluster (*J_FB−FB_* < *J_CM−FB_*). This choice of parameter allowed us to qualitatively reproduce the patterns observed in experiments (see Figure 2 and 5.

**Table 1.**
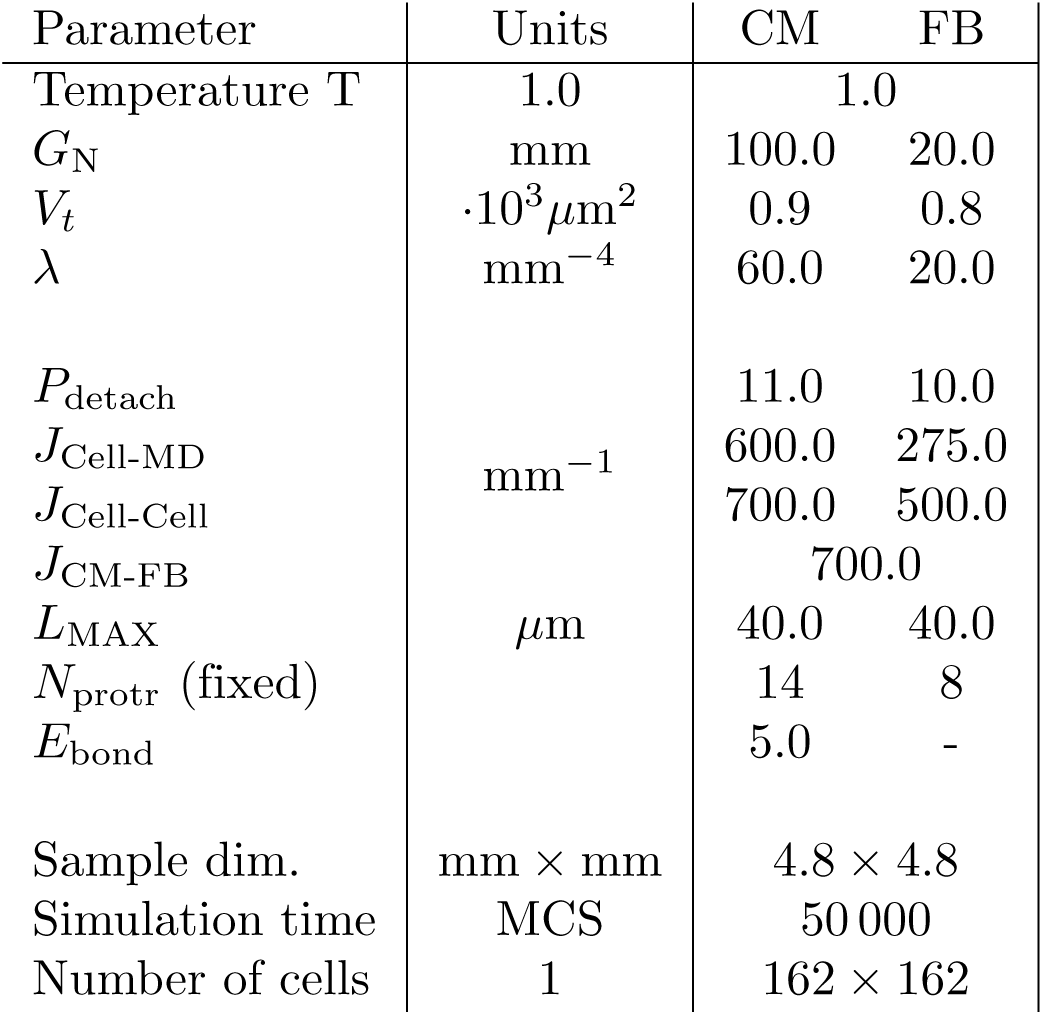
Parameters of the morphological model used in stimulations. CM—cardiomyocytes, FB—fibroblasts.

### 3.2 Simulations of wave propagation

The virtual samples generated with the Cellular Potts Model were then used as a map for electrophysiological studies. The methodology was already described in detail in our previous paper [15]. The parameters of the electrophysiological model were the same as in [15], except for the coupling coefficients that were slightly adjusted (*D_in_* = 0.4 *cm*^2^*/s*, *D_L_* = 0.4 × 10^−2^ *cm*^2^*/s*, *D_T_* = 0 *cm*^2^/*s*) to reproduce the maximal conduction velocity in the control samples with low fibrosis. The same coefficients were then used for all simulations.

## Supporting information

**S1 Video. Raw optical recording of the wave propagation in a neonatal rat cardiac monolayers with with 66% of fibroblasts.** This video is also available at https://youtu.be/3aDmsT1pl3Y

**S2 Video. Growth of the branching pattern.** The left part shows cardiomyocites in red, and fibroblasts in cyan, and the right part highlights connected clusters with different colours. The red cluster is the one that was growing from the center and eventually wired the whole sample. Video can be also accessed at https://youtu.be/s9V86BFcMQY.

**S3 Video. Raw output of the simulation of the wave propagation in a sample with 70% of fibroblasts.** This video is also available at https://youtu.be/lw4p7cen0u4.

**S4 Video. Spontaneous formation of the structure, that produces a uni-directional block, which results in reentry formation.** The virtual sample had 29% of cardiomyocytes. Left pannels show wave propagation from bottom to top, and right panel in the opposite direction. Two bottom row shows enlarged insets of the place, where where a uni-directional block has occurred. The video is also available at https://youtu.be/6LZorTUcJdk.

**S5 Video. Visualisation of spontaneous symmetry breakup and polarization of the cells.** The video is also available at https://youtu.be/RatnaeS7N2M

## Author contributions statement

K.A. conceived the idea of the experiments, A.N. and V.Ts. conducted the experiments, N.K. performed simulations under supervision of A.P. A.N., V.Ts. and N.K. analysed the results. All authors reviewed the manuscript.

## Footnotes

1 http://www.worthingtonbiochem.com/NCIS/default.html

2 http://www.abcam.com/protocols/immunocytochemistry-immunofluorescence-protocol

